# B.1.526 SARS-CoV-2 variants identified in New York City are neutralized by vaccine-elicited and therapeutic monoclonal antibodies

**DOI:** 10.1101/2021.03.24.436620

**Authors:** Hao Zhou, Belinda M. Dcosta, Marie I. Samanovic, Mark J. Mulligan, Nathaniel R. Landau, Takuya Tada

## Abstract

DNA sequence analysis recently identified the novel SARS-CoV-2 variant B.1.526 that is spreading at an alarming rate in the New York City area. Two versions of the variant were identified, both with the prevalent D614G mutation in the spike protein together with four novel point mutations and with an E484K or S477N mutation in the receptor binding domain, raising concerns of possible resistance to vaccine-elicited and therapeutic antibodies. We report that convalescent sera and vaccine-elicited antibodies retain full neutralizing titer against the S477N B.1.526 variant and neutralize the E484K version with a modest 3.5-fold decrease in titer as compared to D614G. The E484K version was neutralized with a 12-fold decrease in titer by the REGN10933 monoclonal antibody but the combination cocktail with REGN10987 was fully active. The findings suggest that current vaccines and therapeutic monoclonal antibodies will remain protective against the B.1.526 variants. The findings further support the value of wide-spread vaccination.

## Introduction

Severe Acute Respiratory Syndrome Coronavirus 2 (SARS-CoV-2) is a highly transmissible and pathogenic coronavirus that became an ongoing pandemic late in 2019. Infection rates have begun to fall, at least in part, due to large-scale vaccination efforts. In addition, treatment of infected patients with monoclonal antibodies against the spike protein have been found to reduce hospitalization and mortality [1, 2]. The recent emergence of SARS-CoV-2 variants raises concerns both with regard to vaccine efficacy and the effectiveness of monoclonal antibody therapy. The vast majority of sequenced SARS-CoV-2 isolates contain a D614G mutation in the spike protein [3] that increases viral infectivity and transmissibility [4–6] and subsequently, variants with multiple mutations in the spike protein and enhanced transmissibility have emerged in the United Kingdom [7–9], South Africa [10], Brazil [11] and the United States [12, 13] raising concerns of diminished neutralization by immune sera-elicited antibodies and escape from therapeutic monoclonal antibodies.

Recent reports have identified a novel variant in New York City termed B.1.526 that was rapidly spreading [14–16]. The variant was identified in November, 2020; by January, 2021 the variant accounted for 5% of genomes sequenced from individuals in New York and by mid-February was detected with a frequency of 12.3% [14–16]. The variant contains several mutations in the spike protein, some of which have not been found in previous variants. Two versions of B.1.526 were identified; both with the D614G mutation and A701V, in addition, the mutations L5F, T95I and D253G which are not present in previously reported variants. One version of B.1.526 contains the E484K mutation which is present in the B.1.351 and B.1.1.248 variant spike proteins and allows for partial escape from immune sera neutralization [17–23]; the other lacks the E484K mutation but has a nearby S477N mutation, which lies within the receptor binding domain (RBD) and thus may influence affinity for the entry receptor ACE2. The D253G mutation is located in the amino-terminal supersite that serves as a binding site for neutralizing antibodies while A701V is located adjacent to the furin processing site. The combination of mutations raises concerns that the B.1.526 variant might evade vaccine elicited and therapeutic antibodies.

Previous studies have shown that the E484K in the B.1.351 spike protein leads to a degree of resistance to neutralization by both infection- and vaccine-elicited antibodies as well as to the REGN10933 therapeutic monoclonal antibody [24–27]. Moreover, the B.1.351 variant spike protein has been found to reduce the level of protection provided by vaccination in populations in which the variant has become prevalent [24, 28].

In this study, we determined the susceptibility of the B.1.526 variants to neutralization by convalescent sera and sera from individuals vaccinated with the Pfizer BNT162b2 [29] and Moderna mRNA1273 vaccines [30] and by Regeneron therapeutic monoclonal antibodies [25, 31]. We found that the B.1.526 variant (S477N) was fully susceptible to neutralization while the B.1.526 with the E484K mutation neutralized with a modest (3.5-fold) reduction in titer by convalescent and vaccine elicited antibodies. The B.1.526 spike proteins were readily neutralized by Regeneron antibody cocktail.

## Results

The B.1.526 variant spike proteins contain the D614G mutation, a shared set of novel mutations (L5F, T95I, D253G, and A701V) and either E484K or S477N, both of which lie within the RBD **(Fig. 1A and B)**. To study the B.1.526 spike proteins, we constructed spike protein expression vectors for both B.1.526 versions and used these to produce lentiviral pseudotypes reporter viruses as previously described. Immunoblot analysis showed that both B.1.526 spike proteins were expressed and processed in transfected 293T cells and that both were incorporated into virions at a level comparable to that of the wild-type (D614G) spike protein **(Fig. 1C)**. The infectivity of B.1.526 variant pseudotypes on ACE2.293T was similar to that of wild-type (**Fig. 1D**).

**Figure.1.**
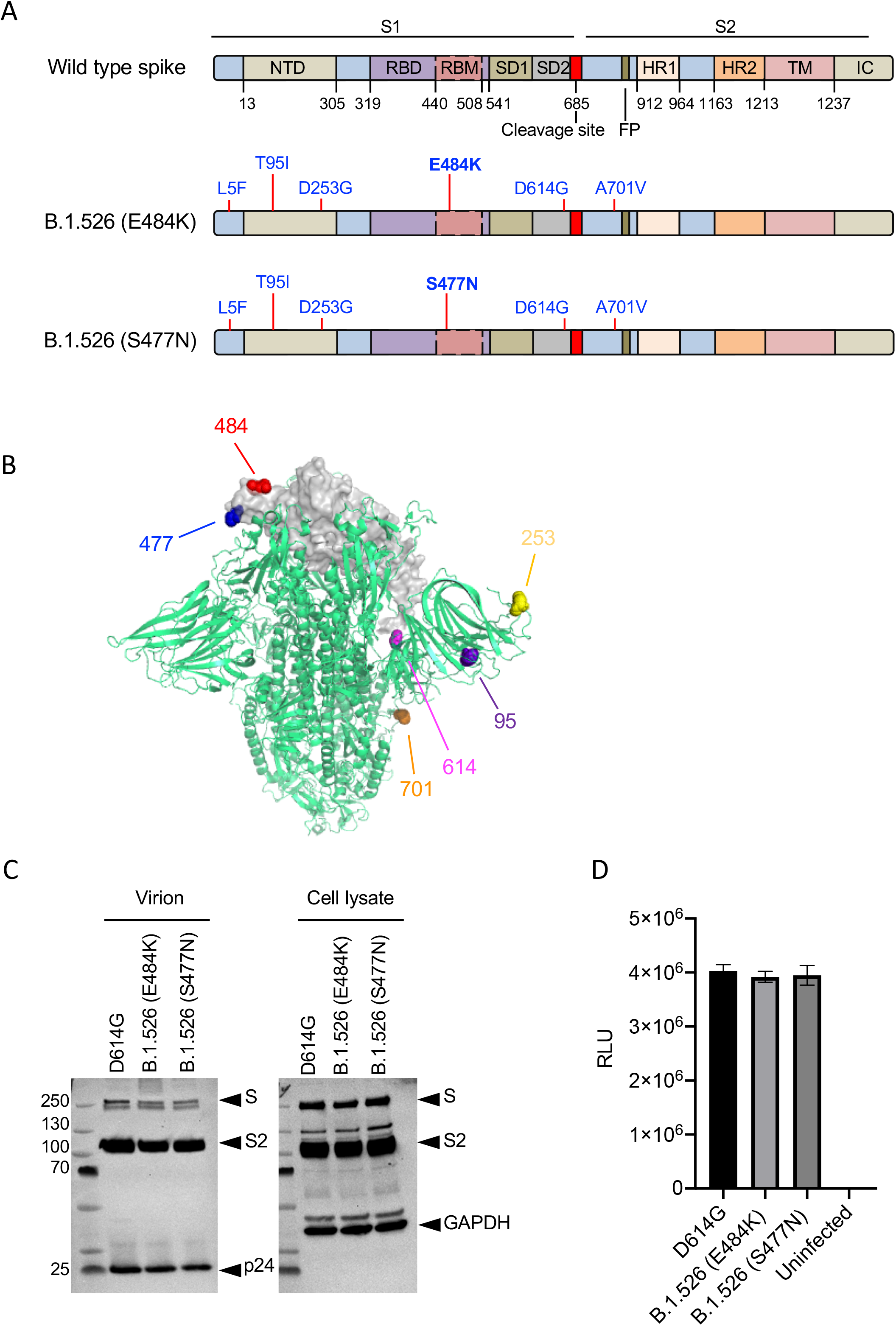
Analysis of B.1.526 pseudotyped lentiviral virions. **(A)** The domain structure of the SARS-CoV-2 spike protein is shown above. NTD, N-terminal domain; RBD, receptor-binding domain; RBM, receptor-binding motif; SD1 subdomain 1; SD2, subdomain 2; FP, fusion peptide; HR1, heptad repeat 1; HR2, heptad repeat 2; TM, transmembrane region; IC, intracellular domain. The location of the mutations in the two B.1.526 spike proteins is diagrammed below with the distinguishing E484K and S477N mutations in bold. **(B)** The location of the B.1.526 variant spike protein mutations are shown on the 3D structure of the trimeric spike protein. One RBD region is shown for simplicity. The 484 (red) and 477 (blue) amino acid residues are indicated. **(C)** Immunoblot analysis of B.1.526 spike protein pseudotyped lentiviral virions and cell lysates of transfected producer cells. The blots were probed for the full-length S and processed S2 proteins and with anti-P24 antibody to detect the virions. GAPDH served as a loading control for the cell lysates. Arrows indicate the full-length spike (S), S2 subunit (S2). **(D)** Infectivity of B.1.526 pseudotyped virus in ACE2.293T cells. ACE2.293T cells were infected with pseudotyped viruses normalized for RT activity. Luciferase activity was measured 2 days post-infection as relative light units (RLU).

To test the ability of convalescent sera to neutralize the B.1.526 viruses, we determined the neutralizing antibody titers of sera from individuals who had been infected prior to April 2020 on viruses with B.1.526, D614G and E484K spike proteins (**Fig. 2A**). The results showed that neutralizing titers against the S477N B.1.526 variant were similar to that of D614G while the neutralizing titers against the E484K B.1.526 variant decreased by 3.8-fold, a modest decrease that was attributed to the E484K mutation.

**Figure.2.**
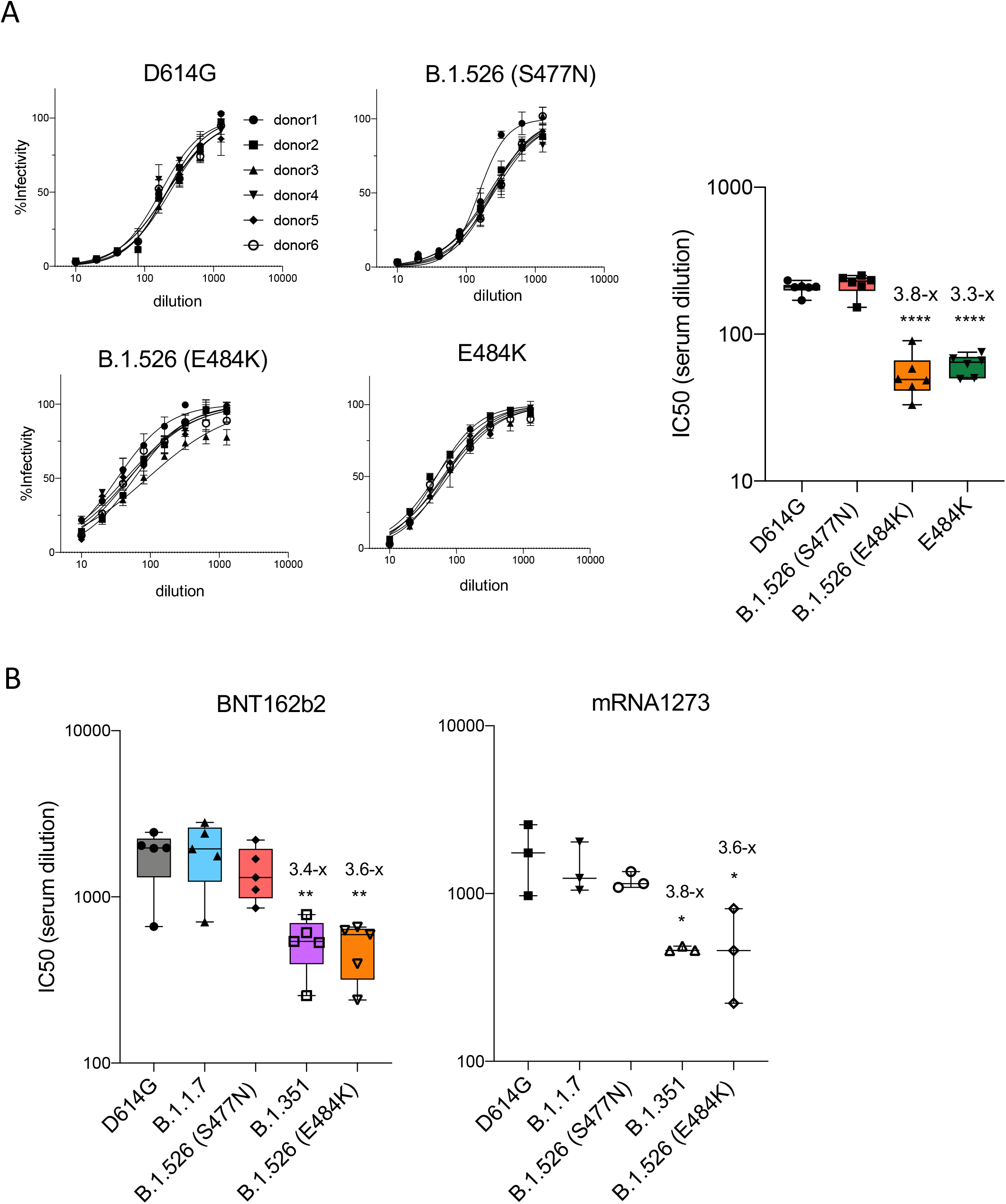
Convalescent serum and antibodies elicited by BNT162b2 and mRNA1273 vaccine neutralize B.1.526 variant spikes. **(A)** Neutralization of viruses with D614G, B.1.526 variant spikes by convalescent sera (n=6). The data are shown as the percentage infectivity in the absence of serum (left). IC50 (serum dilution) of sera from convalescent individuals (n=6) against virus with B.1.526 and E484K variant spikes (right). **(B)** Neutralizing titers of serum samples from BNT162b2 vaccinated individuals (n=5) (left) and mRNA1273 vaccinated donors (n=3) (right) was measured. IC_50_ of neutralization of virus with D614G, B.1.1.7, B.1.351, B.1.526 is shown.

To determine the ability of vaccine-elicited antibodies to neutralize the B.1.526 viruses, we determined neutralizing titers of serum specimens from individuals vaccinated with Pfizer BNT162b2 or Moderna mRNA1273 vaccines. The results showed that BNT162b2 vaccine serum-elicited antibodies neutralized the D614G and B.1.1.7 viruses with similarly high titers while titer for neutralization of B.1.351 was decreased by 3.4-fold (**Fig. 2B left**). Analysis of the B.1.526 titers showed that the S477N version was neutralized with a titer similar to D614G; neutralization titers of the E484K version were decreased by 3.6-fold, a titer similar to that of B.1.351. Neutralization by sera from Moderna vaccine showed a very similar pattern (**Fig. 2B right)**.

Analysis of the Regeneron monoclonal antibodies showed that REGN10987 neutralized both B.1.526 variants with no loss of titer **(Fig. 3A)**. REGN10933 neutralized virus with the S477N B.1.526 spike protein with a high titer but was 12-fold less active against the E484K B.1.526 version **(Fig. 3B)**. The cocktail which forms the REGN-COV2 therapy potently neutralized the B.1.526 spike variants despite the partial loss of neutralizing activity against the E484K version of B.1.526.

**Figure.3.**
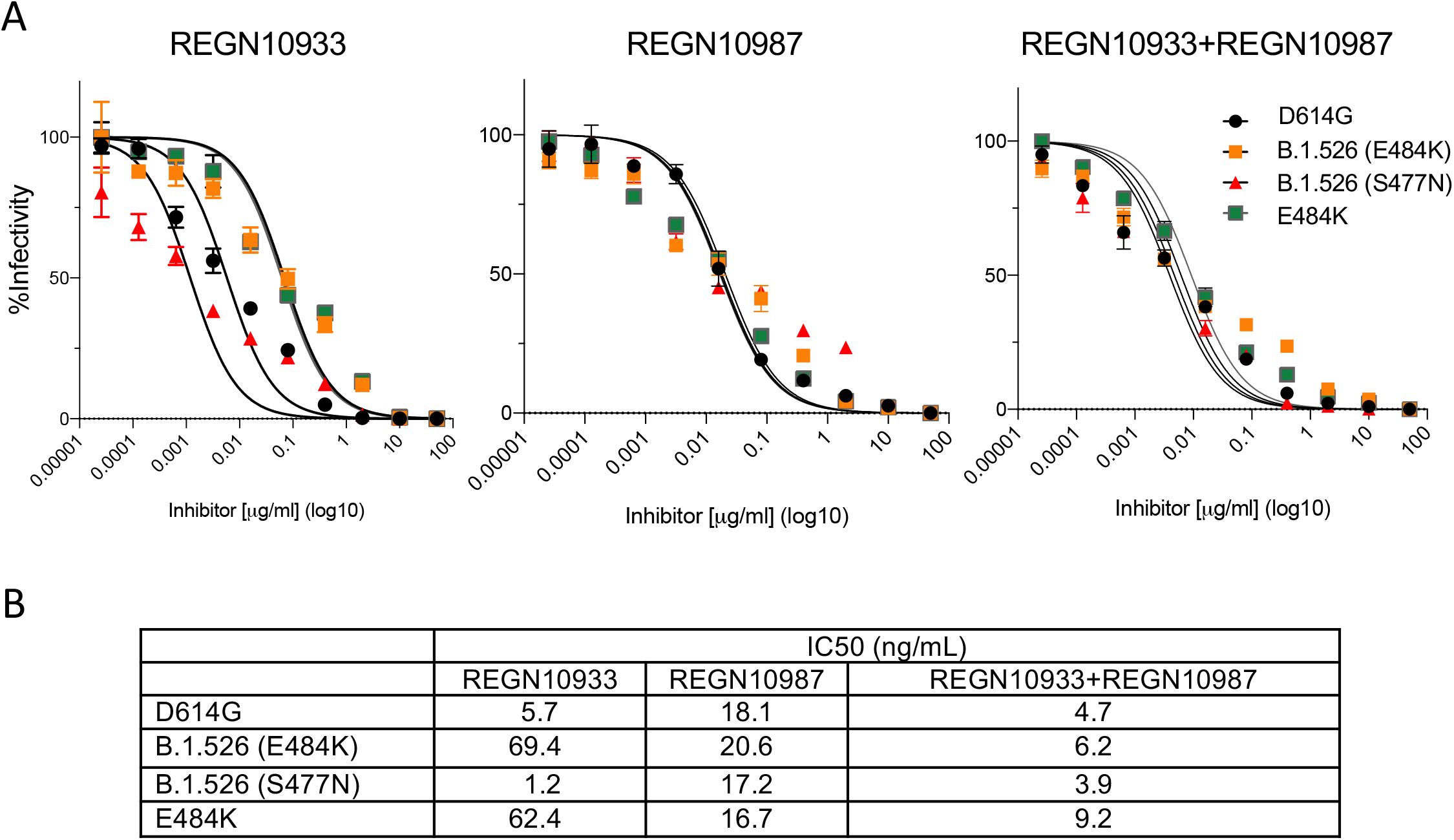
Analysis of B.1.526 neutralization by REGN10933 and REGN10987. Neutralizing titers of REGN10933, REGN10987 antibodies and a 1:1 mixture of the two antibodies was measured on viruses pseudotyped with B.1.526, D614G and E484K spike proteins. **(A)** Neutralization curves of REGN10933, REGN10987 and a 1:1 mixture of REGN10933 and REGN10987 on viruses with the B.1.526, D614G and E484K variant spike proteins. **(B)** IC_50_ of the monoclonal antibodies and the combination cocktail on viruses with the variant spike proteins as calculated using the data in panel A.

## Discussion

The recently identified B.1.526 variant SARS-CoV-2 appears to be increasing in prevalence in New York City raising concerns about reinfection and immunoevasion [14–16]. We report here that both S477N and E484K versions of B.1.526 were neutralized well by convalescent and vaccine-elicited antibodies. The E484K version of B.1.526 did show a significant nearly 4-fold decrease in neutralization by vaccine-elicited antibodies but this represents a modest decrease in titer that is not expected to result in a significant decrease in the protection provided by vaccination and is not expected to result in an increased susceptibility to re-infection. The S477N version was neutralized with no decrease in titer.

Our results showed that REGN10987, which binds to the side of the RBD [25, 31], maintains potent neutralizing activity against both versions of B.1.526 but that REGN10933, which binds to the top face of the RBD that interacts with ACE2, loses 12-fold of its potency against the E484K version. The decrease in neutralizing titer was caused by the E484K mutation and is similar to the previously reported loss of titer against B.1.351 which also bears the mutation [24–28, 32–34]. Despite the partial loss of activity by REGN10933 against B.1.526, the neutralizing activity of the combined antibody cocktail remained high.

Our findings should assuage concerns that the B.1.526 variant will evade protection provided by vaccine-elicited antibodies and suggest that therapeutic antibody therapy will retain its effectiveness against the variant. Nevertheless, B.1.526 appears to be spreading at an alarming rate, demonstrating the value of wide-spread vaccination efforts.

## Methods

### Plasmids

pLenti.GFP.NLuc dual GFP/nanoluciferase lentiviral vector, pcCOV2.Δ19S codon-optimized SARS-CoV-2 spike gene expression vector, HIV-1 Gag/Pol expression vector pMDL and HIV-1 Rev expression vector pRSV.Rev have been previously described [35]. B.1.526 spike mutations were introduced into pcCOV2.Δ19S by overlap extension PCR. All plasmid sequences were confirmed by DNA nucleotide sequencing.

### Cells

293T cells were cultured in Dulbecco’s modified Eagle medium (DMEM) supplemented with 10% fetal bovine serum (FBS) and 1% penicillin/streptomycin (P/S) at 37°C in 5% CO2. ACE2.293T is a clonal cell-line that stably expresses a transfected human ACE2. The cells were maintained in DMEM/1 μg/ml puromycin/10% FBS/1% P/S.

### Human Sera and monoclonal antibodies

Convalescent sera and sera from BNT162b2 or Moderna-vaccinated individuals were collected on day 28 following the second immunization at the NYU vaccine center with written consent under IRB approval (IRB 20-00595 and IRB 18-02037). Donor age and gender were not reported. Regeneron monoclonal antibodies (REGN10933 and REGN10987) were prepared as previously described [36].

### SARS-CoV-2 spike lentiviral pseudotypes

SARS-CoV-2 spike-pseudotyped lentiviruses were produced by co-transfection of 293T cells with pMDL, pLenti.GFP-NLuc, pcCoV2.S-Δ19 (or variant spikes) and pRSV.Rev, as previously described [35]. The viruses were concentrated by ultracentrifugation and normalized by reverse transcriptase RT activity. To quantify neutralizing antibody, sera were serially diluted 2-fold and then incubated with pseudotyped virus (approximately 2.5 × 10^7^ cps) for 30 minutes at room temperature. The mixture was then added to 1 × 10^4^ ACE2.293T cells, corresponding to an MOI of 0.2 in a 96-well cell culture dish. After 2 days, luciferase activity was measured using Nano-Glo luciferase substrate (Nanolight). Luminescence was read in an Envision 2103 microplate luminometer (PerkinElmer).

### Immunoblot analysis

Cells were lysed in buffer containing 50 mM HEPES, 150 mM KCl, 2 mM EDTA, 0.5% NP-40, and protease inhibitor cocktail. Lysates (40μg) were separated by SDS-PAGE and transferred to a polyvinylidene difluoride membrane. The membranes were probed with anti-spike mAb (1A9) (GeneTex), anti-p24 mAb (AG3.0) and anti-GAPDH mAb (Life Technologies) followed by goat anti-mouse HRP-conjugated second antibody (Sigma). The blots were visualized with luminescent substrate (Millipore) and quantified on an iBright CL1000 Imager.

### Quantification and Statistical Analysis

All experiments were performed in technical duplicates or triplicates and data were analyzed using GraphPad Prism 8. Statistical significance was determined by the twotailed, unpaired t-test. Significance was based on two-sided testing and attributed to p< 0.05. Confidence intervals are shown as the mean ± SD or SEM. (*P≤0.05, **P≤0.01, ***P≤ 0.001, ****P≤ 0.0001). The PDB file of D614G SARS-CoV-2 spike protein (7BNM) was downloaded from the Protein Data Bank. 3D view of protein was obtained using PyMOL.

## Acknowledgements

The work was funded by grants from the NIH to N.R.L. (DA046100, AI122390 and AI120898) and to M.J.M. (UM1AI148574), T.T. was supported by the Vilcek/Goldfarb Fellowship Endowment Fund.

## Author contributions

T.T. and N.R.L. designed the experiments. H.Z., T.T. and B.M.D. carried out the experiments and analyzed data. H.Z., T.T. and N.R.L. wrote the manuscript. M.I.S. and M.J.M. provided key reagents and useful insights. All authors provided critical comments on manuscript.

## Competing interests

The authors declare no competing interests.

## Notes

### Competing Interest Statement

The authors have declared no competing interest.

